# Whole-brain modelling supports the use of serotonergic psychedelics for the treatment of disorders of consciousness

**DOI:** 10.1101/2023.12.29.573603

**Authors:** I Mindlin, R Herzog, L Belloli, D Manasova, M Monge-Asensio, J. Vohryzek, A. Escrichs, N Alnagger, P Núñez, ML Kringelbach, G Deco, E Tagliazucchi, L Naccache, B Rohaut, JD Sitt, Y Sanz Perl

## Abstract

Disorders of consciousness (DoC) are a challenging and complex group of neurological conditions characterised by absent or impaired awareness. The current range of therapeutic options for DoC patients is limited, offering few non-invasive pharmacological alternatives. This situation has sprung a growing interest in the development of novel treatments, such as the proposal to study the efficacy of 5HT2A receptor agonists (also known as psychedelics) to restore impaired consciousness. Given the ethical implications of exploring novel compounds in non-communicative individuals, we assessed in silico their effects in the whole-brain dynamics of DoC patients. We embedded the whole-brain activity of patients in a low-dimensional space, and then used this representation to visualise the effects of simulated neuromodulation across a range of receptors representing potential drug targets. Our findings show that activation of serotonergic and opioid receptors shifted brain dynamics of DoC patients towards patterns typically seen in conscious and awake individuals, and that this effect was mediated by the brain-wide density of activated receptors. These results showcase the role of whole-brain models in the discovery of novel pharmacological treatments for neuropsychiatric conditions, while also supporting the feasibility of accelerating the recovery of consciousness with serotonergic psychedelics.

## Introduction

Despite the lack of broad definitions of consciousness as a global brain state, it is accepted that it can be lost or diminished when we fall asleep, or as the effect of drugs, such as those employed to induce general anaesthesia. Consciousness is also impaired in pathological conditions, such as during coma or in post-comatose disorders of consciousness (DoC). The two primary categories within the spectrum of DoC are unresponsive wakefulness syndrome (UWS) that is also coined vegetative state (VS), and minimally conscious state (MCS). UWS is characterised by preserved arousal without any behavioural evidence of consciousness, while MCS represents a condition with richer cognitive processes that are not limited to reflexive processes. Observation of MCS patients demonstrates that, - in contrast with UWS/VS patients -, cortical networks still play a role in their overt behaviour (Giacino et al., 2002; Naccache, 2018). Even if being in a MCS does not necessarily correspond to being in a “minimally conscious” state, it does certainly correspond to a Cortically Mediated State (CMS). DoC are complex neurological diseases with heterogeneous etiologies and presentations, often resulting in a substantial misdiagnosis rate with potentially devastating implications (Fins, 2019).

In the last two decades, the incorporation of neuroimaging techniques resulted in significant improvements regarding the diagnosis and prognosis of DoC patients (Owen, 2020) Data-driven machine learning approaches are capable of identifying brain activity patterns that predict different levels of impaired consciousness, with an accuracy comparable to that of expert neurologists (Comanducci et al., 2020; Hermann et al., 2021; Kondziella et al., 2020; Sitt et al., 2014; Stefan et al., 2018). One downside of these approaches is that they are not based on generative models, therefore they cannot be used to test causal mechanisms underlying the observed differences between groups (Deco & Kringelbach, 2014). In turn, the insufficient understanding of how consciousness is lost in brain-injured patients hinders the development of novel therapeutic interventions (Marino & Whyte, 2022).

Currently, the range of treatments for DoC includes Zolpidem and Amantadine as pharmacological options (Giacino et al., 2012), as well as invasive (Schiff et al., 2007) and non-invasive electrical magnetic stimulation methods, such as transcranial direct current stimulation (tDCS) and transcranial magnetic stimulation (TMS) ((Bourdillon et al., 2019). The pharmacodynamics of the pharmacological interventions are heterogeneous. Zolpidem restores consciousness in chronic patients due to a paradoxical effect (Bomalaski et al., 2017), and its effect wears off along with the drug (Whyte et al., 2014; Whyte & Myers, 2009). In contrast, Amantadine is a N-methyl-D-aspartate receptor antagonist and indirect dopamine agonist capable of inducing both short (5-10 days) and long-term (4-6 weeks) improvements (Rühl et al., 2022), and it was incorporated in DoC practical guidelines for clinicians (Giacino et al., 2018). Unfortunately, available drugs have modest therapeutic efficacy and act via unknown specific mechanisms, thus hindering the development of new options, while clinical trials are difficult to organise due to low number of participants and other limitations intrinsic to patients with conscious impairments (Marino & Whyte, 2022)

In recent years, serotonergic psychedelics have been proposed as a potential new treatment to accelerate the recovery of DoC patients (Scott & Carhart-Harris, 2019). This proposal was not based on pharmacological considerations, but on the measured effects of psychedelics on brain activity, and the interpretation assigned to these changes by theories of consciousness (R. L. Carhart-Harris et al., 2014). Psychedelic drugs are suggestive to increase brain complexity measured using either entropy or Lempel-Ziv complexity (LZC) (M. M. Schartner et al., 2017; Tagliazucchi et al., 2014), both metrics known to decrease during episodes of diminished conscious awareness (deep sleep, general anaesthesia, and DoC) (Koch et al., 2016). At the same time, over the past years the use of psychedelics for treating mental health disorders has shown promising results in conditions such as depression (R. Carhart-Harris et al., 2021; Davis et al., 2021; Goodwin et al., 2022; Palhano-Fontes et al., 2019) obsessive compulsive disorder (Moreno et al., 2006), among others. It is speculated that psychedelics can result in functional rearrangements mediated by serotonin 2A (5-HT2A) receptor agonism, which modulates the emergent whole-brain dynamics, generating principal psychoactive effects of these drugs (Ly et al., 2018). Importantly, the density of this receptor is maximal in key sub-cortical and high-level cortical association areas implicated with conscious information processing (Beliveau et al., 2017; Guldenmund et al., 2012). Taken as a whole, these findings support the proposal by Scott and Carhart-Harris of exploring classic psychedelics as new avenues for the treatment of DoC.

The current consensus is that psychedelic drugs are very safe substances when consumed by healthy individuals under adequate conditions (Nichols, 2016). However, a deeper analysis reveals distinct ethical concerns for the use of psychedelics in non-communicative patients (Peterson et al., 2019). Computational modelling offers an alternative to evaluate the therapeutic action of serotonergic psychedelics without facing the ethical challenges of human *in vivo* experimentation. While this research cannot replace proper clinical trials, it can be used to support the feasibility of this intervention, adding to evidence from other sources, such as animal experimentation.

Here we proposed a framework combining deep learning and computational whole-brain modelling to provide mechanistic understanding of possible pharmacological treatments for DoC, including serotonergic treatments, in a controlled and ethical manner. We used a biophysically-grounded model based on coupled mean-fields representing the dynamics of brain regions at the macroscale, composed by excitatory and inhibitory neural populations (Deco et al., 2014). The strength of connection between nodes is informed through anatomical structural connectome informed by diffusion MRI (dMRI). Using this Dynamic Mean Field (DMF) model, we simulated agonism at multiple receptors by altering the nonlinear response of neuronal populations to synaptic input, weighted by the local density of receptors informed by Positron Emission Tomography (PET) data. This integrative multimodal approach in computational modelling has already demonstrated its effectiveness in capturing the intricate interplay between coupled whole-brain networks and neurotransmitter systems (Kringelbach et al., 2020). As in previous work, we trained a variational autoencoder (VAE) (Kingma & Welling, 2013; Perl et al., 2023) for model visualisation and validation, interpreting the effects of simulated interventions as trajectories in the low-dimensional latent space, and allowing their characterization in terms of well-established complexity metrics sensitive to the level of conscious awareness.

## Results

### Methodological overview

The procedure followed in this work is represented in **Fig 1**. First, we implemented a biologically realistic DMF model (Deco et al., 2014) fitted to fMRI data from DoC patients and healthy controls. The DMF models brain dynamics as the emergent activity of local recurrent excitatory/inhibitory connections, with long-range excitatory connections between cortical regions, scaled by the global connectivity parameter (G). Local inhibition is controlled by the Feedback Inhibitory Current (FIC), which offsets inter-areal excitation and clamps the average excitatory firing rate around 3 Hz. The G parameter, and thus indirectly the FIC, are tuned to maximise the similarity to the empirical brain functional connectivity (FC), as computed from the fMRI Blood-Oxygen-Level-Dependent (BOLD) signals. Finally, the expression of neurotransmitter receptors on each region is obtained from PET maps, and the susceptibility of each region to the activation of these receptors is controlled via a global scaling parameter. Since every region has its own level of receptor expression, this global scaling parameter will introduce regionally heterogeneous neuromodulation.

**Fig 1.**
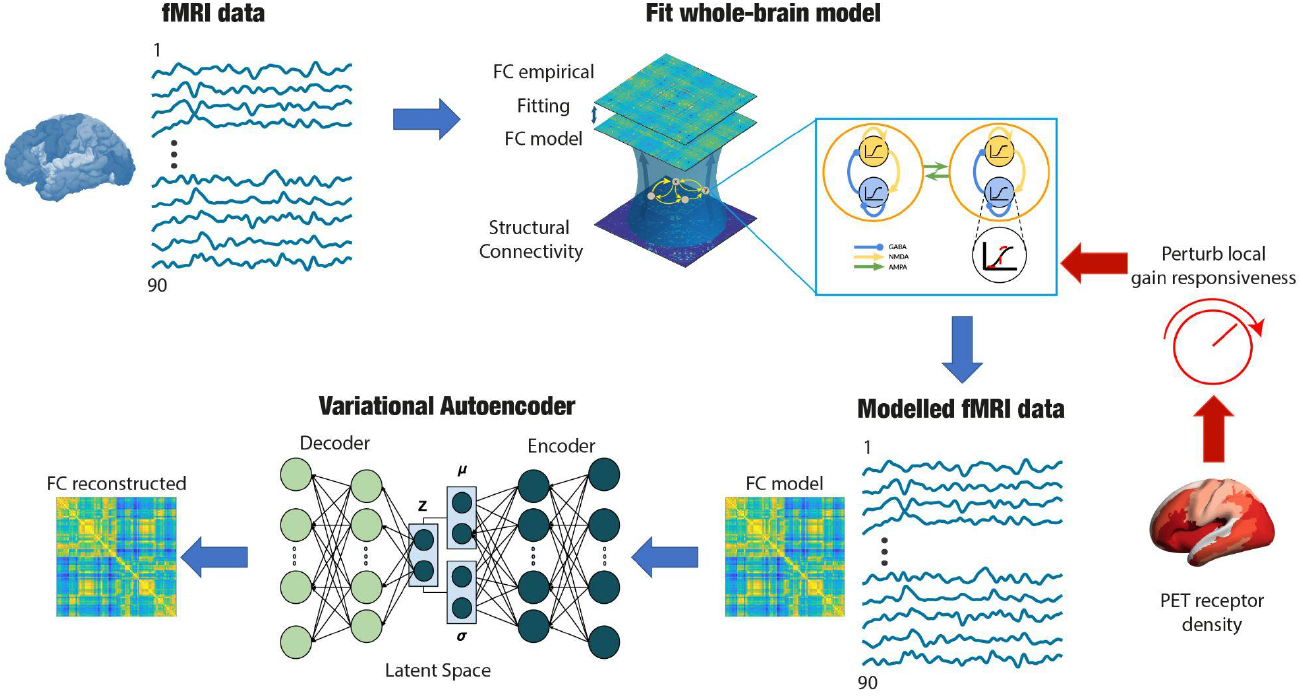
Overview of the whole brain model, variational autoencoder and perturbation pipeline. A biophysical model simulates the local dynamics of a 90 region parcellation, with structural connectivity provided by DTI data. Model parameters are adjusted to fit FC matrices derived from fMRI data. Pharmacological perturbations are simulated by changing the synaptic scaling of each region weighted by PET receptor maps, which provide a measure of local receptor density. Subsequently, a variational autoencoder (VAE) is trained to reconstruct the input data, resulting in a low-dimensional latent space which facilitates the visualisation of the perturbations, while also providing a geometric interpretation of their effect.

After tuning the model, we used the inferred parameters for each condition (*G* and FIC) to generate surrogate FC matrices as a data synthesis procedure (Perl, Pallavicini, et al., 2020) to train a variational autoencoder (VAE), resulting in a two-dimensional encoding of each FC matrix. We then characterised this low-dimensional space in terms of different metrics sensitive to changes in consciousness, such as the Lempev-Ziv complexity, and finally explored the effects of pharmacological interventions. As a result, changes in receptor scalings yield the potential to reflect global shifts in neurotransmission signalling and can explain whole-brain dynamics in response to a drug intervention.

We implemented this procedure with 8 different receptors (serotonergic 5HT2A, 5HT1A, 5HT6; dopaminergic D1, D2; dopamine transporter (DAT); opiate μ, Histamine H3). We selected serotonergic receptors to test the effect of psychedelic drugs, whose main target is the 5HT2A receptor (Nichols, 2016), while the choice of dopaminergic receptors stemmed from their relevance to current treatments for DoC (e.g. Amantadine). Including opiate and histamine receptors completed the spectrum of potential stimulation targets, resulting in a comprehensive array of diverse receptors for analysis.

As a final step, we encoded the pharmacological intervention in the two-dimensional latent space, parametrized by the scaling that modulates the receptor density maps. This combined framework allowed us to represent the interventions as trajectories in the latent space and compute geometrical properties that characterised the intervention, such as distance to the healthy controls, the distance from the original non-perturbed model, as well as changes in brain complexity, network integration, and correlation with the anatomical connectivity.

### Baseline model fitting

To explore the effect of perturbing via different receptors, we first created a *baseline* model for controls and each patient group, namely UWS and MCS. For each value of G, we conducted repeated simulations for the number of subjects in each group and then also repeated the training process 20 times to optimise the SSIM between simulation and our target data at group level. The SSIM is a similarity measure that balances correlation and euclidean distance between the inputs (Wang et al., 2004), in this case the simulated and the empirical FC. The range of G values were defined such that the model always remained in a biologically plausible regime, below hyper-excited and hyper-correlated activity. In **Fig 2A**, the plots demonstrate the fitting performances for each group. In all three groups, a point of instability is reached, resulting in the breakdown of the goodness of fit. Prior to this instance, all the models attain their best fit. We repeated this exploration of G values 15 times. Then, we calculated the optimal value of G for each condition by taking the mean optimal G obtained across iterations (mean+-std: *G*_UWS_*=*1.59+-0.04; *G*_*MCS*_*=*1.75+-0.07; *G*_*CNT*_=2.1+-0.06). The low variability in the results across iterations highlights the robustness of our models. These G values determine how we model each condition with the DMF at the baseline.

**Fig 2.**
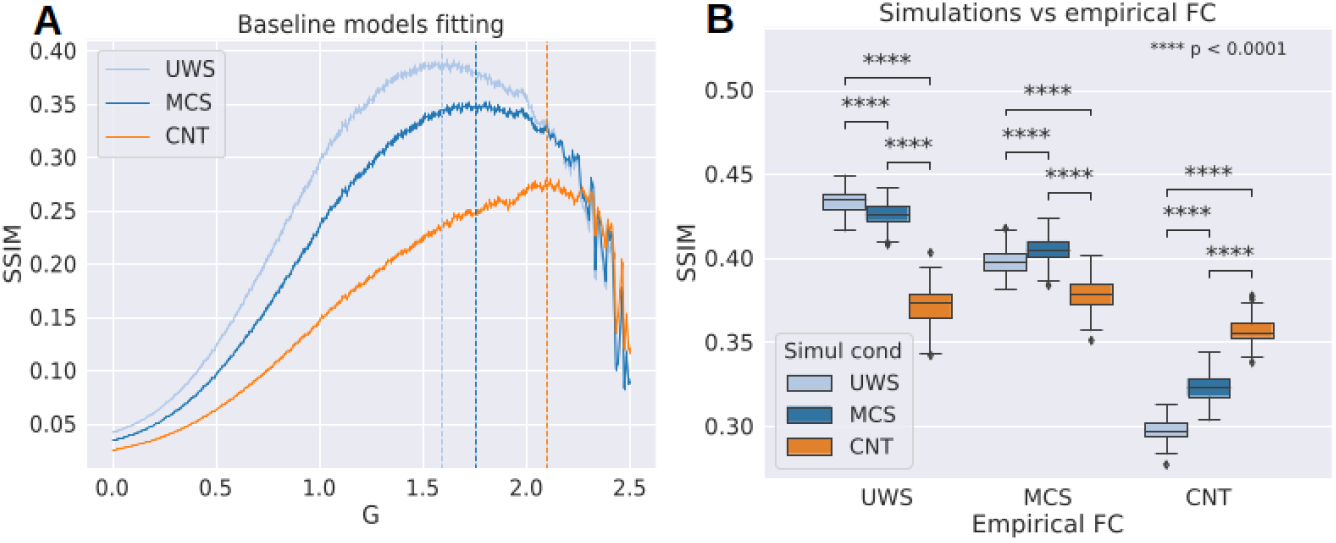
Model fitting and validation of the baseline models. **(A)** Model fit curves where the parameter G is optimised to maximise the SSIM to the empirical FC. The optimal values of G for each condition (mean+-std: GUWS=1.59+-0.04; GMCS=1.75+-0.07; GCNT=2.1+-0.06) align with existing literature, where lower *G* is associated with reduced consciousness. **(B)** The optimal model for each condition exhibits significantly better SSIM values to its corresponding empirical dataset compared to the rest.

As a next step, we assessed if these models were distinguishable between levels of consciousness. To this end we generated 3000 simulations and calculated the SSIM between their FC and the empirical FC of each condition. We performed a Kruskal-Wallis test where we compared the SSIM values obtained between simulations and empirical FCs for each condition. The models simulations were significantly different across states (Empirical UWS / Simulations CNTvsMCS Cohen’s d=6.09 MCSvsUWS Cohen’s d=1.14 CNTvsUWS Cohen’s d=6.83 Empirical MCS / CNTvsMCS Cohen’s d=3.24 CNTvsUWS Cohen’s d=2.35 MCSvsUWS Cohen’s d=0.96 Empirical CNT / MCSvsUWS Cohen’s d=3.51 CNTvsMCS Cohen’s d=−4.67 CNTvsUWS Cohen’s d=−8.47). Additionally, each model had the highest similarity with the condition they were fitted to (SSIM_*uws*_=0.43, SSIM_*MCS*_=0.40, SSIM_*CNT*_=0.35) as shown in **Fig 2B**. As the *G* parameter was decreased, regions did not interact between themselves, reducing integration of the network and stabilising the overall dynamics to those determined by the anatomical structure. This situation was seen for unconscious brain states, whereas the opposite could be seen for conscious controls. Therefore the resulting optimal values of G found for each condition are consistent with this observation (*G*_UWS_*<G*_*MCS*_*<G*_*CNT*_).

### Latent space embedding of baseline models

After fitting the baseline models, we employed 3000 simulations from each condition to train a variational autoencoder (VAE) for reconstructing the input data, following previous work (Perl et al., 2023). The VAE is a type of generative model that combines the strengths of traditional autoencoders with the probabilistic framework of variational inference. The variational component allows the model to learn the statistical distribution of the data in the latent space rather than just mapping the data to a fixed point. By learning the statistical distribution of the input data, the VAE constructs a continuous and smooth latent space (Kingma & Welling, 2013), called *manifold*, that captures the underlying structure of the simulated FC. This results in a meaningful representation of the data, where similar inputs are mapped closer together and dissimilar inputs are mapped farther apart in the latent space. The use of the 2-dimensional latent space allowed us to visualise a *perturbational landscape*, summarising the impact of all possible pharmacological perturbations applied to the DoC models.

Once the VAE was trained, we encoded a test set to observe how the different conditions organised in the latent space, as depicted in **Fig 3A**. The different conditions formed separate clusters with very small overlap, which means that conditions were distinguishable within the latent space embedding. Additionally, there appeared an ordering along an axis starting from lower consciousness (UWS), passing through an intermediate stage (MCS), and finally to the fully conscious state (CNT). **Fig 3B** quantifies this placement as the distribution of the euclidean distance for the DoC simulations with respect to the CNT centroid (||UWS||=2.93+-0.51; ||MCS||=1.64+-0.45). The difference of means between these distributions had a very large effect size (Cohens’d=2.66). This distribution of distances allowed us to determine how much dynamics were changed with respect to our healthy target. Aside from identifying the regions where simulations of baseline conditions are projected we want to gain interpretability of what it means in terms of consciousness for a FC matrix to be projected in any given point of the latent space. This is presented through the combination of panels in **Fig 3C-E**. The highlighted area shows the value at the centroid of each condition. As seen in **Fig 3C**, the embedded FC matrices from the control group appear in regions that exhibit higher LZ complexity than the corresponding DoC regions (Median,IQR LZC_UWS_=0.024,0.001, LZC_MCS_=0.028, 0.02, LZC_CNT_=0.031,0.001 UWSvsMCS Cohen’s d=−2.69 MCSvsCNT Cohen’s d=−2.85 UWSvsCNT Cohen’s d=−5.12). To calculate the LZ complexity on the FC matrices, we traversed all the columns within a single row, following the methodology established in previous studies (Varley et al., 2020). Integration was measured by the average shortest path length between regions (Neudorf et al., 2023) with lower values expressing more efficiency in information exchange (Median,IQR INT_UWS_=1.30, 0.15, INT_MCS_=1.17, 0.10, INT_CNT_=1.06, 0.06 UWSvsMCS Cohen’s d=1.33 MCSvsCNT Cohen’s d=1.70 UWSvsCNT Cohen’s d=2.88). As shown by previous research, loss of consciousness abolishes functional interactions that are not directly mediated by structural links, thus making FC patterns of DoC patients more similar to the underlying structural connectivity (SC) pattern (Barttfeld et al., 2015; Demertzi et al., 2019). **Fig 3E** shows these values for our three conditions (Median,IQR SC-CORR_UWS_=0.70, 0.02, SC-CORR_MCS_=0.67, 0.01, SC-CORR_CNT_=0.64,0.015 UWSvsMCS Cohen’s d=2.08 MCSvsCNT Cohen’s d=3.47 UWSvsCNT Cohen’s d=5.25)

**Fig 3.**
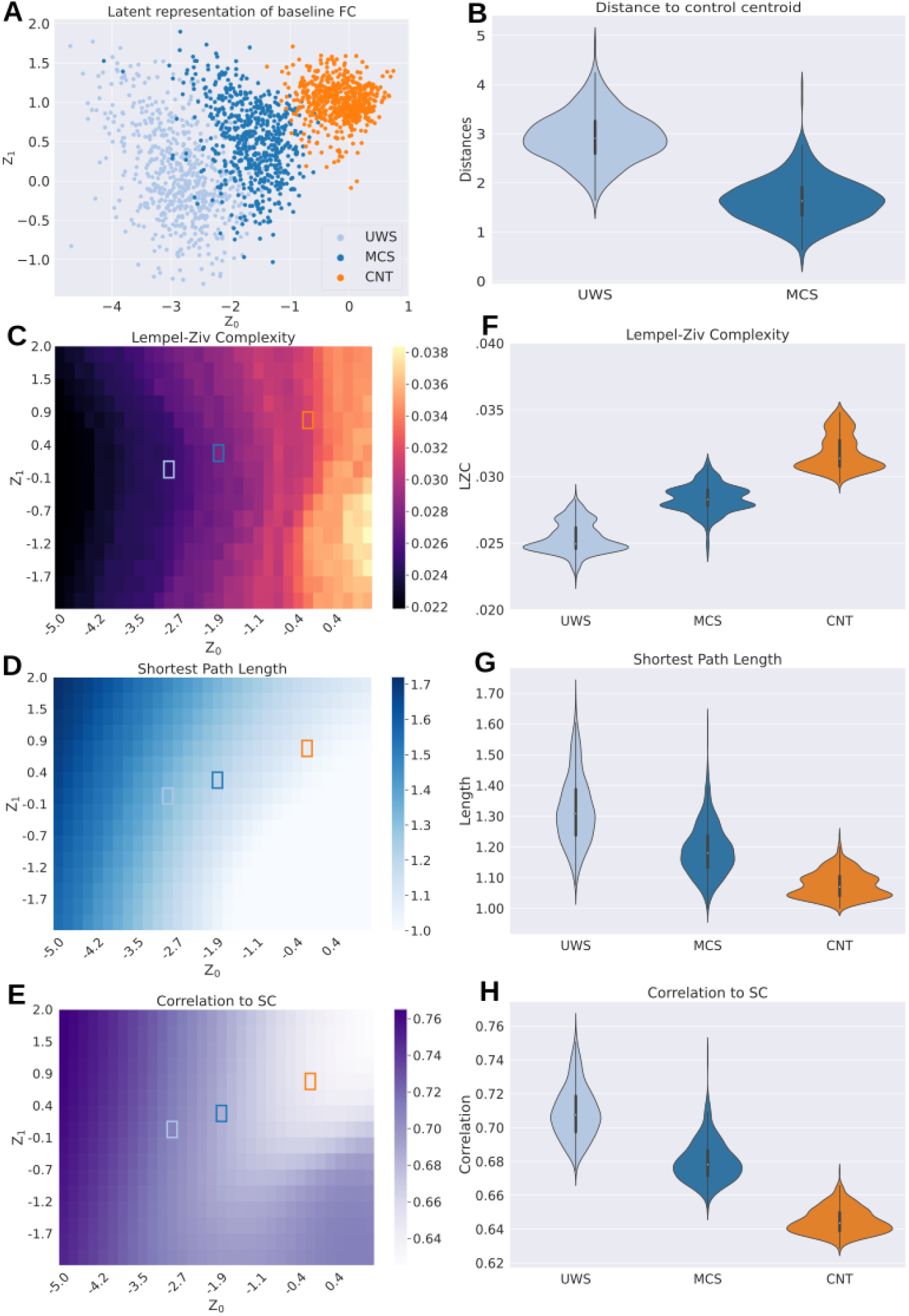
Groups are distinguished and characterised by their latent space embedding in an ordered manner. **(A)** Scatter plot of FC matrices obtained from the three baseline models for the three groups of participants. The groups are clearly separated within the embedded regions. Embedded FC matrices from Disorders of Consciousness (DoC) conditions are distinguished by **(B)** their distance from the centroid of the FC matrices from the healthy control (CNT) condition, which is the largest for unresponsive wakefulness syndrome (UWS) and smallest for minimally conscious state (MCS). We sampled FC matrices from a grid to calculate different metrics over the latent space: **(C)** LZ complexity, **(D)** shortest path length (a proxy for integration), and **(E)** correlation to structural connectivity. Each square corresponds to a point from that grid and has therefore an associated value. Highlighted squares indicate the position of the centroids of each group in this space. The values of these metrics in the regions corresponding to each condition are consistent with their interpretation concerning consciousness. **(F-G-H)** Distribution of the previous measures when sampling the embedded points from panel **A**.

### Pharmacological perturbations produce transitions away from baseline dynamics

Once we obtained an interpretable latent space of our baseline simulations, we simulated pharmacological perturbations of the DoC models using the PET density maps of different receptors. As mentioned before, these maps give a density value for each node that is then scaled by the *scaling* parameter (see Methods). The perturbation consists of increasing this value in a range between 0 (no receptor intervention) to 1 with a step of 0.01. Embeddings of the perturbations applied to MCS and UWS groups are shown in **Fig 4A-B**, respectively. For each receptor map we embedded in the latent space the simulated FC after applying the scaling value in a given step. The resulting trajectory comes from encoding the simulated FC matrices for all steps in the scaling range. Different receptor trajectories reach different distances and the proximity to the control centroid is limited by its starting point (**Fig 4C-D**). Another interesting observation is that all the trajectories seem to move approximately in the same direction, i.e. in parallel to the line that unites the three groups from states of more diminished consciousness towards the group of wakeful conscious controls.

**Fig 4.**
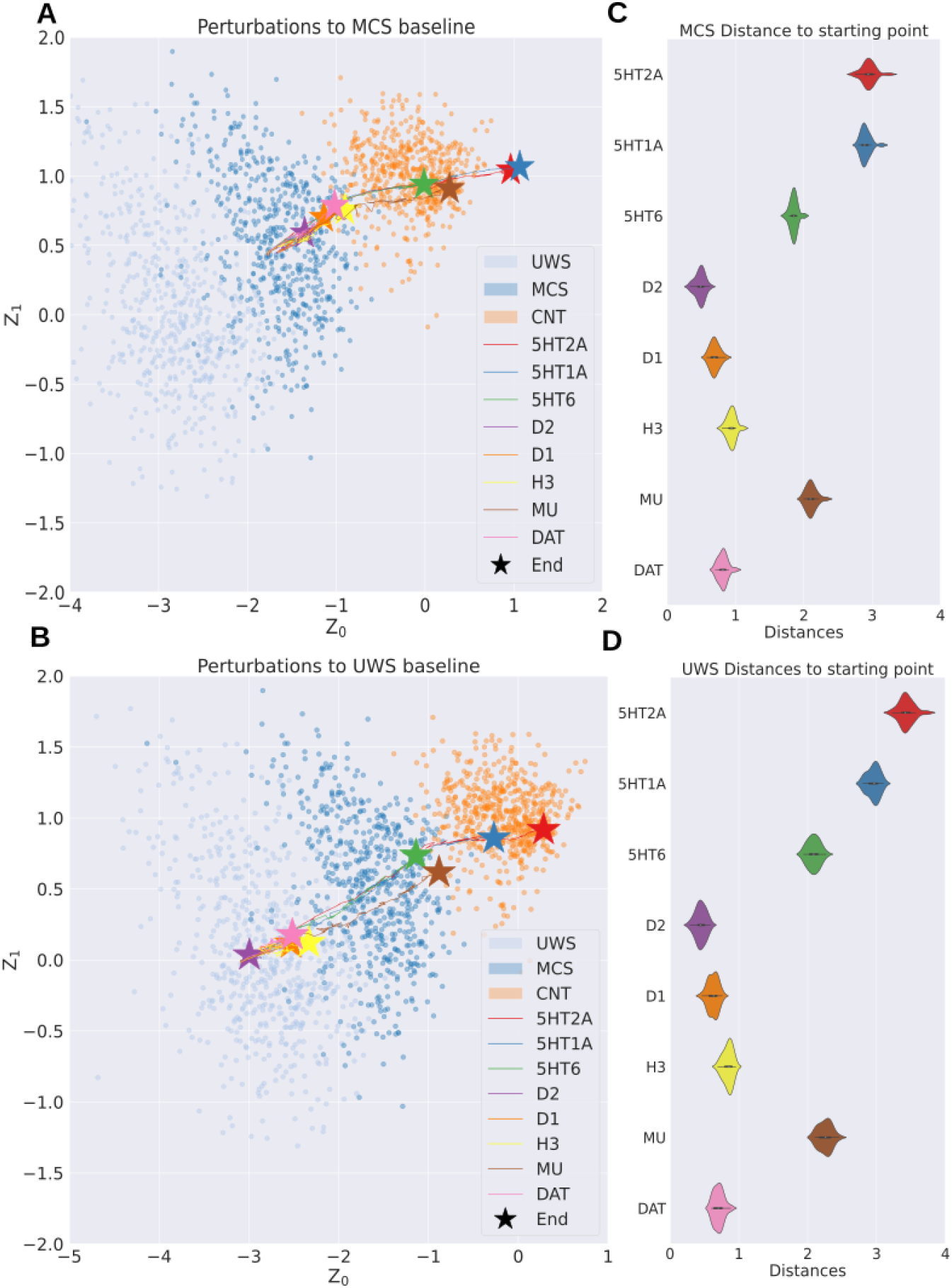
Hierarchy of target neurotransmitter receptors based on how their simulated activation displaces whole-brain dynamics towards conscious wakefulness. Panels **(A)** and **(B)** display embedded trajectories for perturbing the MCS and UWS baseline models, respectively, at the different receptors. All perturbations displace the embedded FC matrices along the same axis in latent space, indicating significant variations in the final distance covered across the entire perturbation range. This is presented in panels **(C)** and **(D)**, where the covered distances are similar, with differences in the maximum distance reached. Specifically, for UWS, the perturbations did not deviate as far from their starting point, as was observed for the MCS models. Simulated perturbations at the serotonergic receptors and the MU receptor presented the most significant impact on the models.

These results show that despite exhibiting different anatomical heterogeneities, the perturbed dynamics exist only in certain regions in latent space, while others are forbidden, thus narrowing the repertoire of possible transitions. The main difference between the trajectories is the distance from the end point to the starting point (Longest and shortest distance achieved for each condition: Max-MCS_5HT1A_=3.49, Max-UWS_5HT2A_=4.61, Min-MCS_D2_=1.62, Min-UWS_D2_=0.74).

### Relationship between distribution of receptors and perturbation trajectories

We investigated the relationship between the final distance reached by the trajectory and two attributes of the receptor maps: the mean density of the map and a structural divergence (SD) metric (see Methods). Briefly, the (SD) is the euclidean distance between the density map and the node strength, which represents the average structural connection strength from each node to the rest. A low SD implies that the density map and the node strength are similar, suggesting that the receptor has high density values in strongly connected nodes, or hubs, and vice versa.

We found a strong correlation between the mean density of the map and the final distance of the trajectory (**Fig 5A-B**, MCS Pearson correlation: 0.89, p=0.003; UWS Pearson correlation: 0.96, p=0.0001). The difference in correlation values can be attributed to the curvature of the trajectory after crossing the CNT region in MCS perturbations, which leads to changes in linear distance from the starting point. Furthermore, **Figs 5C-D** demonstrate that SD is not correlated with the length of the trajectory. The combination of these results suggests that the neuromodulation of a given receptor displaces the dynamics of a baseline DoC model independently of the topology of the stimulated nodes. Instead, it primarily depends on the receptor density throughout the brain.

**Fig 5.**
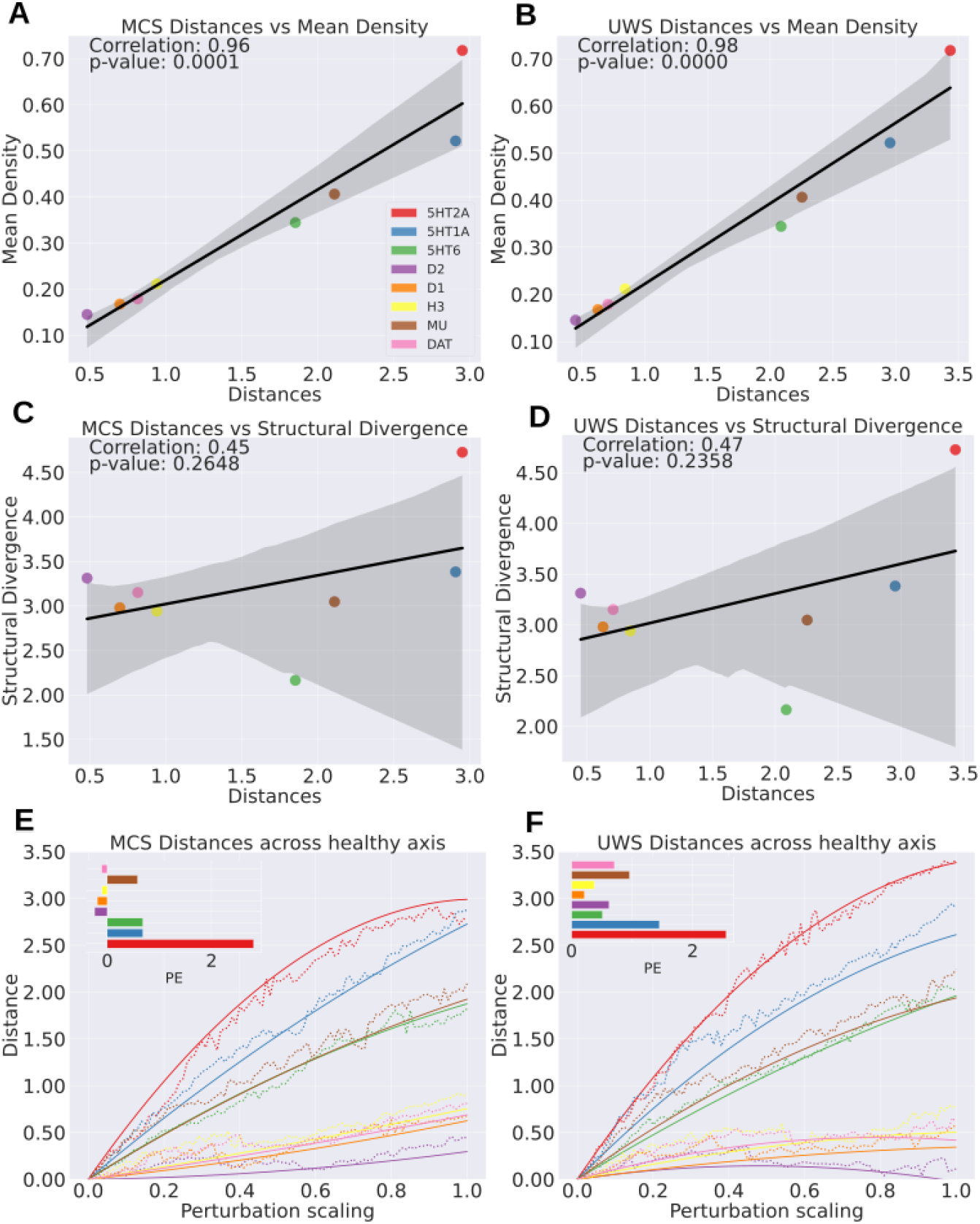
Geometric properties show that perturbation trajectories are related to the density distribution of stimulated receptors. Panels **(A)** and **(B)** show that the maximum distance reached from the starting point exhibits a strong linear correlation with the mean receptor density. In panels **(C)** and **(D)**, in contrast, we show that there is no significant correlation between the maximum distance reached and the “structural divergence” (representing the spatial relationship between receptor maps and the weight vector of connections). This finding highlights the extent to which the model dynamics can be shifted independently of the spatial arrangement of receptors. **(E-F)** Dotted lines show the perturbational distance accumulated as the *scaling* increases. Solid lines indicate the quadratic regression fitted to the cumulative distances. The inset shows the “perturbational efficiency” (PE) for each receptor. The 5HT2A receptor has the highest PE despite reaching a similar overall perturbational distance than the 5HT1A.

Additionally, we assessed how intensely each receptor should be stimulated to shift brain dynamics. Different drugs may require different relative dosages to alter brain dynamics depending on their pharmacokinetic and pharmacodynamic properties. Ideally, good candidates for treatments should exert their action with small doses to avoid undesired side-effects due to the stimulation of off-target receptors. For this purpose, we estimated how quickly the perturbation could displace the model towards CNT-like dynamics. We calculated the dot product between each projected simulation and an axis drawn between the starting point of the models and the CNT centroid (**Fig 5E-F)**. Trajectories may reach similar distances along this axis, yet this can occur at different *speeds*. To quantify this, we applied a quadratic regression fit to the cumulative distance obtained as the *scaling* increases. The concavity of this fit expresses the rate at which the distance is achieved. Therefore, we label “perturbational efficiency” (PE) the negative quadratic coefficient of the best fit, which represents the aforementioned rate. We observe that 5HT2A has the highest PE, even though it has a similar perturbational distance than the other serotonin receptor subtype, 5HT1A.

## Discussion

DoC encompasses a set of devastating conditions, affecting not only the lives of the patients themselves but also causing emotional distress among their families and caretakers. The diagnosis of DoC patients is very challenging, also resulting in a difficult evaluation of potential treatment options. This and other issues, such as heterogeneous etiologies and small available populations, hinder the systematic exploration of new pharmacological alternatives. Currently used drugs became available as a consequence of indirect assumptions, instead of being optimised specifically for the treatment of DoC patients. The purpose of this study is to demonstrate the feasibility of using biophysically-grounded whole-brain models to inform the effect of targeted receptor stimulation in the clinical status of DoC patients. We were able to map different pharmacological interventions into two-dimensional space trajectories, from which we derived features indicative of treatment feasibility and efficacy. Moreover, successful perturbations not only recovered conscious-like dynamics, but also restored to baseline levels of the three metrics previously introduced for the objective quantification of conscious awareness.

Our results highlight that stimulation of two serotonin receptor subtypes (2A and 1A) restores dynamics indicative of conscious wakefulness. Interpreted within the context of previous proposals to investigate psychedelics as treatment options in DoC (Scott & Carhart-Harris, 2019), this result can be taken as an additional suggestive element favouring further steps, such as the exploration of animal models. Interestingly, even though the 5HT2A receptor is the main target of psychedelic drugs (i.e. all psychedelics are full or partial agonists at this receptor), we also found comparable results for the 5HT1A receptor. Psychedelic compounds are divided into two main families depending on their chemical structure: substituted tryptamines (including N,N-Dimethyltryptamine and psilocybin) and substituted phenethylamines (such as mescaline). Of these families, tryptamines present significant agonism at serotonin 5HT1A/1A receptors (Nichols, 2016). This raises the possibility that the potential therapeutic effect of psychedelics on DoC patients is not universal for all psychedelics, but can change from compound to compound. It is also important to note that not all 5HT2A receptor agonists induce psychedelic effects (López-Giménez & González-Maeso, 2018), thus it could be possible that a non-psychedelic serotonergic agonist is eventually capable of improving the clinical status of DoC patients.

Our model also predicted that stimulation of opioid µ receptors should have a positive impact on DoC patients. This result is consistent with a recent publication demonstrating that use of opioid drugs during surgery correlates with consciousness improvement (Ge et al., 2023). A similar role of µ receptor stimulation is suggested by the study of neurorehabilitation and recovery of brain-injured patients after severe COVID-19 infections, since for MCS patients the amount of oxycodone (an opioid analgesic drug) received during treatment correlated with their recovery indices after the infection (Gurin et al., 2022). It has been speculated that opioid compounds could have neuroprotective properties, which could be beneficial for brain injured patients (Vaidya et al., 2018). These studies should be taken with care, however, as they report observational findings instead of the results of clinical trials designed to evaluate the safety and efficacy of opioids in DoC patients.

The validity of the perturbational analysis depends on the capacity of the baseline biophysical models to capture the relevant features of whole-brain dynamics. The results of the model fitting are consistent with previous studies, such as the work of Lopez-Gonzales and colleagues, showing that a less biophysically-realistic whole-brain model optimises the reproduction of fMRI DoC patient dynamics with low coupling (G) values (López-González et al., 2021). The same behaviour was demonstrated using the same model fitted to statistical observables derived from the turbulent brain dynamics regime (Escrichs et al., 2022). In our case, the optimal G values correlated with the expected level of consciousness per group (i.e. UWS < < MCS < CNT), in accordance with previous work using this model (Luppi, Mediano, et al., 2023). Interestingly, this behaviour seems to be invariant with respect to the choice of model and its optimisation targets, thus being indicative of a neurobiological principle.

Another key point of our analysis is the data-driven synthesis of training samples for the VAE. The use of data synthesis or augmentation techniques is a standard procedure when training deep learning algorithms; in this case, data synthesis was obtained as the output of a generative dynamical system trained to capture the whole-brain FC of participants (Perl, Pallavicini, et al., 2020; Tubaro & Mindlin, 2019). In addition, using the output of the whole-brain model to train the VAE provided a dataset with fixed underlying parameters and controlled initial conditions, which is difficult to obtain from empirical recordings. While the VAE was not strictly necessary to fit the data and simulate the perturbations, it provided a framework to facilitate the interpretation and visualisation of the results, which is necessary considering the high-dimensional nature of fMRI data (Perl, Bocaccio, et al., 2020) endowed each point in latent space with the value of relevant metrics metrics, namely LZC, integration, and correlation to structure (Barttfeld et al., 2015; Crone et al., 2014; Demertzi et al., 2019; Tononi & Edelman, 1998) Thus, the construction of the latent space allowed us to demonstrate that the three groups (UWS, MCS and CNT) appeared organised within a one dimensional direction representing the extent of consciousness impairment, which is aligned with previous results obtained using phenomenological models (Perl et al., 2023). As expected, this direction was also characterised by a decrease in whole-brain LZ complexity and integration, as well as an increase in correlation to anatomical connectivity, of the sampled FC matrices, affirming that the chosen metrics are sensitive to the level of impaired consciousness of the patients.

Previous research investigated the capacity of simulated external perturbations (e.g. TMS and tDCS) to transition towards states of increased consciousness (Perl et al., 2021, 2023) Here, we adopted biophysical models capable of representing the effects of pharmacological perturbations, resulting in a first exploration of simulated drug treatment in DoC patients. In combination with receptor density PET maps of different neuromodulatory systems (Hansen et al., 2022), we introduced heterogeneities in the model by changing the slope of the synaptic scaling function, weighted by local receptor density estimates (Deco et al., 2018; Herzog et al., 2023). It is important to note that the *scaling* allows us to understand large-scale neurotransmission in terms of the regional heterogeneity of susceptibility to endogenous and exogenous modulatory signals. Consequently, changes in the *scaling* yield the potential to reflect global shifts in neurotransmission signalling and explain whole-brain dynamics in response to a drug intervention, but these changes are not directly modelling the dosage of a drug. For each value of *scaling* we projected the generated FC into our latent space. As the *scaling* increased, the projected simulations moved from the lower consciousness regions to that corresponding to the healthy controls in a continuous manner. Leveraging the VAE latent space representation, these trajectories could be quantified using simple geometrical properties, as well as by the features previously assigned to each point of the space. We found that perturbing some receptors induced a significant shift in the dynamics of the DoC model as determined by the travelled distance as the *scaling* parameter increased. Of note, the least effective in terms of reaching a maximum change in the dynamics, were the dopaminergic receptors, which are currently used currently as treatment options (Giacino et al., 2012; Rühl et al., 2022). This discrepancy could arise due to downstream effects of dopaminergic stimulation not included in our model, or due to the fact that dopaminergic drugs used to treat DoC are non-specific with regards to the targeted dopamine receptor subtype. Future work should attempt to characterise the effect of multi-target stimulation in DoC patients.

Our results show that the mean density of the receptor is highly correlated with the maximum distance reached after the perturbation; however, this result was not found for the local node strength, suggesting that the underlying topological structure has a limited effect on the outcome of the perturbation. Additionally, we measured the impact of different receptors by determining how much scaling was required to shift brain dynamics towards a healthy state. Even though some receptors eventually reached the same overall perturbational distance, they did so at different speeds. Notably, the 5HT2A receptor, the main target of serotonergic psychedelics, presented the highest efficiency in this regard.

The present work contributes to demonstrating the usefulness of *in silico* experimentation to investigate perturbations to transition between different brain states. It has been recently stated that this type of modelling approach can be a crucial step towards personalised treatments. (Kringelbach et al., 2020; Luppi, Cabral, et al., 2023). Leveraging the concept of *digital twins* (Vohryzek et al., 2023) and the idea of *phase 0 clinical trials* (Dagnino et al., 2023; Luppi, Cabral, et al., 2023) these computational models offer unique opportunities to explore and predict the effects of therapeutic interventions. Through virtual simulations of patient-specific brain dynamics, “digital twins” provide a testing ground for different treatment strategies, opening doors for personalised and targeted approaches to manage DoC. Additionally, *phase 0 clinical trials* present a platform for early exploratory studies, allowing researchers to assess the safety and efficacy of potential treatments in a limited patient cohort before scaling up to larger trials.

### Limitations

A main limitation of our work is the lack of interaction between neuromodulatory systems in the model. Future work should address this with a more sophisticated modelling of the neuromodulation, for example by perturbing multiple receptors from the same neuromodulatory system. For instance, in the case of LSD, despite its main action being exerted via the serotonergic system, it also exhibits agonism for dopaminergic receptors (Marona-Lewicka & Nichols, 2007). Other large scale data-driven works have shown that most mind-altering drugs can be understood in terms of contributions from multi-neurotransmitter systems (Ballentine et al., 2022; Luppi, Hansen, et al., 2023). In the same direction, the impact of the heterogeneities in the model should be explored via different parameters depending on the drug that is modelled. For instance, sedative compounds such as ketamine and memantine are postulated to exert their effects by modulating the balance between excitatory and inhibitory mechanisms. This hypothesis can be experimentally evaluated by manipulating the FIC parameter.

Additionally, there are interesting directions to take with respect to non pharmacological perturbations such as central thalamic Deep Brain Stimulation, which requires implanting electrodes, or repetitive transcranial magnetic stimulation, non invasive. Although there are theoretical benefits for them, the clinical results are not conclusive or still in early stages (Vanhoecke & Hariz, 2017; Zhang et al., 2021) which could be benefited from *in silico* experimentation.

Another limitation of our work is the lack of structural connectivity included in the DoC patient model construction. The aberrant function presented by the brains of DoC patients is ultimately underwritten by structural damage. As our patient models estimate the structural connectivity from a set of healthy controls, this assumption omits key aspects of simulating patient brain function and resulting perturbations. Moreover, utilising individual DTI data could also serve to address the matter of personalisation. As stated in by Luppi and colleagues (Luppi, Cabral, et al., 2023), given the heterogeneity of DoC it is imperative to create models that are fitted individually. Future publications should aim to improve the accuracy of such models via incorporating patient structural connectivity in their generation. An alternative would be to fit the effective connectivity (Hahn et al., 2019) at an individual level to obtain an indirect estimate of the structural connectivity of the subject. Beyond the particular case of our work, we believe it would be an interesting step to create a multi-modal biophysical model to profit from abundant EEG data acquired from DoC patients. This type of data has been proven useful to diagnose DoC patients (Hermann et al., 2021; Sitt et al., 2014) and could contribute improved temporal resolution which is lacking from fMRI data.

### Conclusions

We provided *in silico* support for the use of serotonergic compounds in the treatment of DoC patients, while also introducing a computational framework to investigate the effect of pharmacological stimulation in neuropsychiatric conditions. Drug development is a lengthy and expensive process, and only a small percentage of the assayed compounds are eventually adopted in clinical practice. The drugs that are currently used to treat DoC patients were not developed for this objective, instead, they were repurposed after their efficacy was discovered. It must be emphasised that our method cannot replace the evidence generated by clinical trials. However, adding whole-brain biophysical models to the arsenal of methods available for the discovery of novel pharmacological treatments could facilitate the concentration of efforts on promising leads, even in cases where the ethical implications could outweigh the unknown benefits to the patients, as is the case of serotonergic psychedelics.

## Methods

### Patients

Our study encompassed a total of 11 patients in MCS (with 5 females; mean age ± s.d., 47.25 ± 20.76 years), 10 patients in UWS (with 4 females; mean age ± s.d., 39.25 ± 16.30 years), and 13 healthy controls (including seven females; mean age ± s.d., 42.54 ± 13.64 years), as detailed in (Escrichs et al., 2022). The clinical evaluation and Coma Recovery Scale - Revised (CRS-R) scoring (Kalmar & Giacino, 2005) were conducted by trained medical professionals to ascertain the participants’ levels of consciousness. A diagnosis of MCS was assigned to patients displaying behaviours potentially indicative of awareness, such as visual tracking, reaction to pain, or consistent response to commands. Conversely, patients were categorised as UWS if they exhibited arousal (eye-opening) without any indications of awareness, never displaying purposeful voluntary movements. This study received approval from the local ethics committee Comité de Protection des Personnes Ile de France 1 (Paris, France) under the designation ‘Recherche en soins courants’ (NEURODOC protocol, no. 2013-A01385-40). Informed consent was obtained from the relatives of the patients, and all investigations adhered to the principles of the Declaration of Helsinki and the regulations of France.

### Anatomical connectivity

*We employed diffusion MRI (dMRI) data from a cohort of 16 healthy right-handed participants (5 women, mean age: 24*.*8 ± 2*.*5) gathered at Aarhus University, Denmark. The research protocol was approved by the internal research board at CFIN, Aarhus University, and received ethical clearance from the Research Ethics Committee of the Central Denmark Region (De Videnskabsetiske Komiteer for Region Midtjylland). All participants provided written informed consent before participating in the study*.

### Anatomical connectivity acquisition

The imaging data was acquired during a single session using a 3 T Siemens Skyra scanner at CFIN, Aarhus University. The structural MRI T1 scan employed the following parameters: voxel size of 1 mm^3^; reconstructed matrix size of 256 x 256; echo time (TE) of 3.8 ms, and repetition time (TR) of 2300 ms.

For the dMRI data collection, a TR of 9000 ms, TE of 84 ms, flip angle of 90 degrees, reconstructed matrix size of 106 x 106, voxel size of 1.98 x 1.98 mm with a slice thickness of 2 mm, and a bandwidth of 1745 Hz/Px were used. The dataset was recorded with 62 optimal nonlinear diffusion gradient directions at b = 1500 s/mm^2^. Additionally, one non-diffusion weighted image (b = 0) was acquired per every 10 diffusion-weighted images. The acquisition of dMRI images employed both anterior-to-posterior phase encoding direction: one in anterior to posterior and the opposite in the rest.

### Resting-state fMRI acquisition

MRI images were acquired using a 3 T General Electric Signa System. T2*-weighted whole-brain resting-state images were obtained through a gradient-echo EPI sequence with axial orientation (200 volumes, 48 slices, slice thickness: 3 mm, TR/TE: 2400 ms/30 ms, voxel size: 3.4375 × 3.4375 × 3.4375 mm, flip angle: 90°, FOV: 220 mm^2^). Additionally, an anatomical volume was generated utilising a T1-weighted MPRAGE sequence during the same acquisition session (154 slices, slice thickness: 1.2 mm, TR/TE: 7.112 ms/3.084 ms, voxel size: 1 × 1 × 1 mm, flip angle: 15°).

### Probabilistic Tractography analysis for anatomical data

In this study, we utilised previously obtained structural connectivity data from (Deco et al., 2017).Briefly, the averaged whole-brain structural connectivity matrix involved a three-step process: first, defining regions based on the AAL template used in functional MRI data; second, estimating connections (edges) between nodes using probabilistic tractography within the whole-brain network; third, averaging data across participants. In brief, the FSL toolbox’s linear registration tool (www.fmrib.ox.ac.uk/fsl, FMRIB, Oxford) (Jenkinson et al., 2002) was used to co-registrer the EPI image with the T1-weighted structural image. The T1-weighted image was co-registered to the T1 template of ICBM152 in MNI space (Collins et al., 1994). Combining and reversing these transformations, we applied them to map the AAL template (Tzourio-Mazoyer et al., 2002) from MNI space to the EPI native space, preserving labelling values through nearest-neighbour interpolation for accurate brain parcellations in individual native space. For further details please refer to the original work (Deco et al., 2017)

### Resting state pre-processing and FC estimation

Resting state data preprocessing was conducted using FSL (http://fsl.fmrib.ox.ac.uk/fsl), following the methodology outlined (Escrichs et al., 2022). In summary, resting state fMRI data were processed using MELODIC (multivariate exploratory linear optimised decomposition into independent components) (Beckmann & Smith, 2004). The process involved discarding the initial five volumes, motion correction using MCFLIRT (Jenkinson et al., 2002), brain extraction using the brain extraction tool (BET) (Smith, 2002), application of spatial smoothing with a 5 mm FWHM Gaussian kernel, rigid-body registration, high pass filtering with a cutoff of 0.01Hz, and application of single-session independent component analysis (ICA) with automatic dimensionality estimation. Subsequently, for patients, lesion-driven artefacts were identified and regressed out, along with noise components, independently for each subject, utilising FIX (FMRIB’s ICA-based X-noiseifier) (Griffanti et al., 2014). Lastly, FSL tools were employed to co-register the images and extract the time series from the AAL atlas (Tzourio-Mazoyer et al., 2002) for each subject in MNI space, encompassing 90 cortical brain regions. Subsequently, the mean time series for each parcellated region were extracted, followed by the calculation of Pearson correlations between the time series of each pair of regional areas to derive the interregional FC matrices.

### Parcellation

Drawing from our earlier comprehensive whole-brain investigations, we adopted the AAL atlas, specifically focusing on the 90 cortical and subcortical non-cerebellar brain regions (Tzourio-Mazoyer et al., 2002). This atlas served as the foundation for integrating all structural, functional, and neuromodulation data. Leveraging FSL tools, we harnessed the publicly accessible receptor density maps in MNI space to derive the mean receptor density for each distinct AAL region in every individual.

### Whole brain modelling

In this study, we employed a network model to simulate spontaneous brain activity at the whole-brain level, where individual brain areas were represented as nodes, and the white matter connections between them were depicted as links. The dynamic mean field (DMF) model, as proposed in (Deco et al., 2014), was utilised to describe the activity in each brain area. This DMF model offers a reductionist approach, summarising the activity of interconnected excitatory (E) and inhibitory (I) spiking neurons into a simplified set of dynamical equations. The model is based on the earlier work of (Wong & Wang, 2006). Within the DMF framework, excitatory synaptic currents 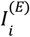 are mediated by NMDA receptors, while inhibitory currents 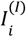 are mediated by GABA receptors.

Each brain area *i* in the DMF model consists of mutually interconnected E and I neuronal pools. Additionally, inter-area coupling between two areas *n* and *p* is exclusively established at the E-to-E level and is adjusted based on the structural connectivity *C*_*ij*_ (for details, refer to Methods - Anatomical Connectivity). This configuration enables the simulation of brain activity and interactions between different regions, providing insights into the dynamics of large-scale neural networks.

More specifically, the DMF model at the whole-brain level is expressed by the following system of coupled differential equations:

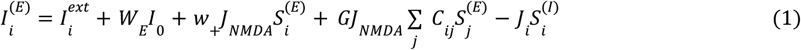

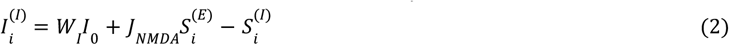

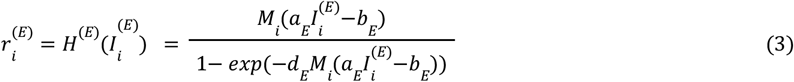

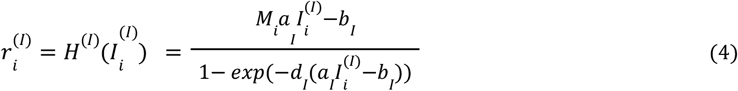

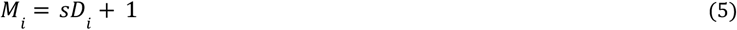

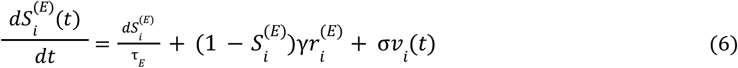

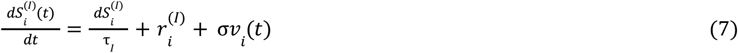

In our computational model, each brain area n was characterised by excitatory (E) and inhibitory (I) pools of neurons, where 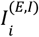 (in nA) represents the total input current, and 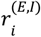 (in Hz) represents the firing rate. Additionally, 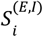 denotes the synaptic gating variable. The neuronal nonlinear response functions, *H*^(*E,I*)^, were applied to convert the total input currents received by the E and I pools into firing rates, 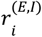, based on the input-output function proposed by (Abbott & Chance, 2005). The model employed specific gain factors, threshold currents, and constants to determine the shape of the curvature of *H* around the threshold.

In this study we used receptor density maps estimated using PET tracer studies obtained by (Hansen et al., 2022). All PET images were registered to the MNI-ICBM 152 nonlinear 2009 (version c, asymmetric) template and subsequently parcellated to the 90 region AAL atlas (Tzourio-Mazoyer et al., 2002). More details on the acquisitions and limitations of this dataset can be found on (Hansen et al., 2022). This processed dataset provided us with a quantitative measure of receptor densities in each AAL region denoted as *D*. These density values were normalised by dividing by the maximum value, ensuring that *max*(*D*_*i*_) = 1. Leveraging *D*_*i*_, we modulated the firing rates 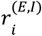 of the excitatory and inhibitory neuronal pools in each brain region, drawing inspiration from (Abbott & Chance, 2005). Receptors were thus used to modulate the gain of the neuronal response function *H*^(*E,I*)^ in each brain area.

The parameters of the DMF model were calibrated to mimic resting-state conditions, ensuring that each isolated node exhibited the typical noisy spontaneous activity with a low firing rate (*r* < 3 Hz) observed in electrophysiology experiments. Additionally, the inhibition weight, *J*_*i*_, was adjusted for each node *i* to achieve Feedback Inhibition Control (FIC). This regulation, described in (Deco et al., 2014), ensured that the average firing rate of the excitatory pools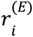 remained clamped at 3 Hz even when receiving excitatory input from connected areas. The FIC was shown to lead to a better prediction of resting-state functional connectivity (FC) and more realistic evoked activity.

The synaptic gating variable for excitatory pools, 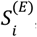, was governed by NMDA receptors, with specific decay time constant tNMDA = 0.1 s and a constant g = 0.641. On the other hand, the average synaptic gating in inhibitory pools depended on GABA receptors with a decay time constant tGABA = 0.01 s. The overall effective external input was represented by *I*_0_ = 0.382 nA, with specific weights *W*_*E*_ = 1 and *W*_*I*_ = 0.7 for excitatory and inhibitory pools, respectively. The model further considered recurrent excitation with a weight of *w*_+_ = 1.4 and excitatory synaptic coupling with weight *J*_*NMDA*_ = 0.15 nA. Gaussian noise, *v*_*i*_, with an amplitude of σ = 0.01 nA was introduced in the system.

In our whole-brain network model, we considered N = 90 brain areas after parcellation of the structural and functional MRI data. Each area *n* received excitatory input from all structurally connected areas p into its excitatory pool, weighted by the connectivity matrix Cnp derived from dMRI data. All inter-area E-to-E connections were equally scaled by a global coupling factor G. This global scaling factor was adjusted to optimise the system’s working point, where the simulated activity best matched the empirical resting-state activity of participants under placebo conditions. To explore different scenarios, simulations were run for a range of G values between 0 and 2.5 with an increment of 0.025 and a time step of 1 ms. Each G value underwent 20 simulations of 192 s duration, mirroring the empirical resting-state scans of 10,11 and 13 participants for the UWS, MCS and CNT conditions respectively.

### Generated BOLD signal

To map the simulated mean field activity to a BOLD signal, we adopted the generalised hemodynamic model proposed by (Stephan et al., 2007). The BOLD signal in each brain area *i* was computed based on the firing rate of the excitatory pools 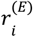. In this model, an increase in the firing rate led to an increase in a vasodilatory signal, *s*_*i*_, which was further subject to auto-regulatory feedback. The changes in blood inflow, *f*_*i*_, were proportionally influenced by this signal, resulting in alterations in blood volume *v*_*i*_ and deoxyhemoglobin content *q*_*i*_. The relationships between these biophysical variables were governed by the following equations:

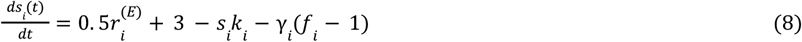

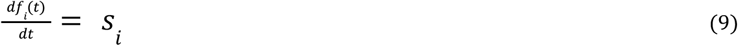

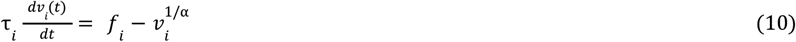

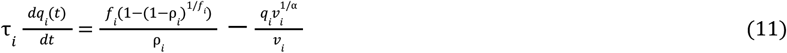

where ρ is the resting oxygen extraction fraction, τ is a time constant and α that represents the resistance of the veins. Finally, the BOLD signal in each area *i, B*_*i*_, is a static nonlinear function of volume, *v*_*i*_, and deoxyhemoglobin content, *q*_*i*_, that comprises a volume-weighted sum of extra- and intravascular signals:

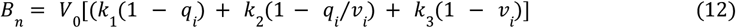

All biophysical parameters were taken as in (Stephan et al., 2007). To concentrate on the frequency range where resting-state activity appears the most functionally relevant, both empirical and simulated BOLD signals were band pass filtered between 0.01 and 0.1 Hz. As with the empirical data we derived the interregional FC matrices between each paris of nodes by calculating the Pearson correlations of their time series.

### Metrics

#### Lempel-Ziv Complexity

Typically, LZC calculations are applied to signals such as EEG data (Dolan et al., 2018). However, in our study focusing on functional connectivity (FC) matrices, we adapted this method by binarizing and flattening these matrices as in previous work (M. Schartner et al., 2015; Varley et al., 2020). This process allowed us to derive a vector from the matrices, facilitating the computation of LZC. Initially, each pixel’s amplitude within the image is assessed, and based on its relation to the median value, it is converted into a binary sequence. The “1”s represent pixels above the median, while the “0”s denote those below it. Subsequently, this binary sequence undergoes a sequential scan to identify distinct structures, thereby constructing a “dictionary of patterns.” The complexity of the flattened image is then evaluated based on the number of patterns present within this dictionary. It’s crucial to note that flattened images with consistent patterns tend to have lower LZ complexity due to a smaller number of distinct patterns, whereas images lacking clear patterns require larger dictionaries, resulting in higher LZ complexity.

#### Integration

To assess the level of integration within the brain networks derived from our FC matrices, we employed the NetworkX library in Python. NetworkX provided the necessary tools to compute the Average Shortest Path Length (ASPL) across these networks. This serves as a proxy measure for integration within complex networks. It quantifies the average minimum number of steps required for information or signals to travel between all pairs of nodes in the network. In our study, the nodes represented brain regions, and the edges denoted the functional connections between these regions. The calculation of average shortest path length involved traversing the network graph and computing the shortest path between all possible pairs of nodes. Let *V* be a set of nodes in *G*, the conversion of FC matrix to graph, *d*(*s, t*) the shortest path from *s* to *t*, and *n* is the number of nodes in *G*

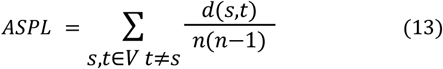

#### Variational Autoencoders

We employed a Variational Autoencoder (VAE) (Kingma & Welling, 2013) to encode the functional connectivity matrices *C*_*ij*_ into a low-dimensional representation. VAEs are neural networks that map inputs to probability distributions in a latent space, allowing for regularisation during training to generate meaningful outputs after decoding the latent space coordinates. The VAE architecture (depicted in **Fig 1**) comprises three components: the encoder network, the middle variational layer, and the decoder network.

The encoder network is a deep neural network utilising rectified linear units (ReLU) as activation functions and includes two dense layers. This part of the network bottlenecks into the two-dimensional variational layer, represented by units z_1_ and z_2_, which span the latent space. The encoder applies a nonlinear transformation to map the functional connectivity matrices (*C*_*ij*_) into Gaussian probability distributions in the latent space. Conversely, the decoder network mirrors the encoder architecture and reconstructs matrices *C*_*i*\**j*_ from samples of these distributions.

The VAE is trained using error backpropagation via gradient descent to minimise a loss function consisting of two terms. The first term is a standard reconstruction error computed from the units in the output layer of the decoder. The second term is a regularisation term measured as the Kullback-Leibler divergence between the distribution in latent space and a standard Gaussian distribution. This regularisation term ensures continuity and completeness in the latent space, ensuring that similar values are decoded into similar outputs and that these outputs represent meaningful combinations of the encoded inputs.

For training the VAE model, we generated 5000 correlation matrices (*C*_*ij*_) corresponding to healthy control, unresponsive wakefulness syndrome (UWS), and minimal conscious state (MCS) conditions. The model’s hyperparameters were optimised using a training set, which was created by randomly splitting the dataset into 80% training and 20% test sets. The training procedure involved using batches with 128 samples and 50 training epochs, with the Adam optimizer and the loss function described in the previous paragraph.

## Acknowledgements

We express gratitude to all the individuals who participated in the studies. Additionally, we extend our thanks to the dedication and support provided by the clinicians at the Neuro ICU, DMU Neurosciences, APHP Sorbonne Université, Hôpital de la Pitié Salpêtrière, Paris, France. Moreover, we acknowledge the invaluable contributions of patient families whose consent and understanding significantly contribute to the advancement of the field. We finally thank the financial support of the FLAG-ERA research funding organisation (project ModelDXConsciousness). Y.S.P is supported by European Union’s Horizon 2020 research and innovation programme under the Marie Sklodowska-Curie grant 896354. M.L.K. is supported by the Center for Music in the Brain, funded by the Danish National Research Foundation (DNRF117), and Centre for Eudaimonia and Human Flourishing at Linacre College funded by the Pettit and Carlsberg Foundations. ET is supported by grants FONCYT-PICT (2019-02294), CONICET-PIP (11220210100800CO), and ANID/FONDECYT Regular (1220995). G.D. was supported by The project NEurological MEchanismS of Injury, and Sleep-like cellular dynamics (NEMESIS) (ref. 101071900) funded by the EU ERC Synergy Horizon Europe; The NODYN Project PID2022-136216NB-I00 financed by the MCIN/AEI/10.13039/501100011033/FEDER, UE., the Ministry of Science and Innovation, the State Research Agency and the European Regional Development Fund; The AGAUR research support grant (ref. 2021 SGR 00917) funded by the Department of Research and Universities of the Generalitat of Catalunya,

## Contributions

**Mindlin I:** Conceptualization, Methodology, Software, Formal analysis, Investigation, Writing - Original Draft, Visualization. **Herzog R:** Software, Conceptualization, Writing - Review & Editing. **Belloli L:** Conceptualization, Writing - Review & Editing. **Manasova D** Software, Conceptualization, Writing - Review & Editing, **Monge-Asensio M:** Software, Resources, Writing - Review & Editing, **Vohryzek J:** Resources, Writing - Review & Editing. **Anira Escrichs:** Resources, Writing - Review & Editing. **Alnagger N:** Writing - Review & Editing. **Núñez Novo P:** Writing - Review & Editing. **Gosseries O:** Writing - Review & Editing, **Morten L. Kringelbach:** Writing - Review & Editing. **Deco G:** Conceptualization, Writing - Review & Editing. **Tagliazucchi E:** Conceptualization, Writing - Review & Editing, **Naccache L:** Resources, Writing - Review & Editing. **Rohaut B:** Resources, Writing - Review & Editing. **Sitt* JD:** Supervision, Conceptualization, Project Administration, Funding Acquisition, Writing - Review & Editing. **Sanz Perl* Y** Supervision, Methodology, Software, Conceptualization, Project Administration, Writing - Review & Editing

